# Biomimetic robots reveal flexible adjustment of sexual signalling in a wild invertebrate

**DOI:** 10.1101/2023.06.11.544488

**Authors:** Joe A. Wilde, Safi K. Darden, Jordan D. A. Hart, Michael N. Weiss, Samuel Ellis, Tim W. Fawcett

## Abstract

Sexual signals are often structured in bouts, which can be adjusted in response to changes in the signaller’s physical and social environment. For example, we might expect individuals to adjust their own signalling behaviour in response to changes in the signalling behaviour of rivals, because this can affect their relative attractiveness to potential mates. In this study, we used a biomimetic robot to experimentally manipulate rival waving behaviour in a wild population of fiddler crabs (*Afruca tangeri*), and investigated whether this leads to changes in the activity and waving behaviour of a focal male. Analysing the focal male’s behaviour using hidden Markov models and linear hurdle models, we found no evidence that the focal male’s waving rate changed in response to changes in the behaviour of the robotic rival. However, bouts of waving lasted longer when the robotic rival was waving at a fast rate. Focal males were also less likely to enter their burrow when the robotic rival was waving, and spent less time in their burrow if they did enter. These results reveal tactical adjustment of behaviour by competing signallers, and highlight the flexible nature of bout-structured sexual displays.

## 1 Introduction

Males from a diverse array of taxa perform courtship displays to attract females for mating opportunities. These include auditory displays such as the Túngara frog’s (*Engystomops pustulosus*) calls or the songs of many passerine birds, visual displays such as the side-blotched lizard’s (*Uta stansburiana*) ‘push-up’ displays or the fiddler crab’s (family *Ocypodidae*) major claw waving, or multi-modal displays such as the greater sage grouse’s (*Centrocercus urophasianus*) ‘strut’ and ‘pop’ or the ‘dance’ and wing-snap of the golden collared manakin (*Manacus vitellinus*). These examples, along with most other examples of courtship displays, are performed in a bout-structure where the courtship display is repeated over a relatively short period of time, leading to temporal clusters of display events (Bateson & Martin, 2021; Perry et al., 2019).

Multiple theories have been suggested to explain why display behaviours are repeated in a bout structure. If the function of a display is to broadcast information about the underlying quality of a signaller and there is error in broadcasting this information via the display (either originating from the performer or the receiver), then repeated displays will reduce this error and give a more accurate representation of signaller quality (Mowles & Ord, 2012). However, display repetition may also act as a way for signallers to demonstrate their quality to potential mates. For example, if a male’s display rate shows his ability to withstand the cost of rapidly producing an already costly display multiple times, signal repetition rate will act as a reliable indicator of signaller quality and potential mates should prefer signallers with higher display rates (Mowles & Ord, 2012). Female preference for faster courtship rates has been demonstrated in wolf spiders (*Shizocosa uetzi*, Shamble et al., 2009; *Hygrolycosa rubrofasciata*, Parri et al., 1997), fiddler crabs (*Austruca mjoebergi*, Callander et al., 2011; Mowles et al., 2018) and crickets (*Gryllus lineaticeps*, Wagner and Reiser, 2000).

It has also been demonstrated that there are aspects of bout-structured sexual signals that signallers tactically adjust in response to their external environment. For example, when a predator is added to their tank, male guppies (*Poecilia reticulata*) will reduce the rate at which they perform visually conspicuous sigmoidal displays (Endler, 1987), but will increase the frequency of these displays in high light levels where these displays are more effective (Chapman et al., 2009). Males of the fiddler crab *Austruca perplexa* have been shown to adjust the wave rate and height of the major claw wave depending on the distance to females (How et al., 2008). Male greater sage grouse (*Centrocercus urophasianus*) adjust the length of their signalling bouts in response to the distance to a female and whether she appears to be inspecting the signalling male or not (Perry et al., 2019). These findings not only highlight the dynamic nature of these displays, raising questions about how such highly dynamic and flexible traits can honestly indicate signaller quality (see Hollon et al., 2023), they also highlight the myriad ways in which bout-structured sexual signals can be adjusted. The interval between each individual display, the speed at which each individual display is performed, the length of signalling bouts, the interval between signalling bouts, the height to which an ornament is raised, and the volume or pitch of an auditory display are just a few examples of the many facets of sexual displays a signaller can tactically adjust in response to external cues (Patricelli et al., 2016).

Given that signallers rarely compete for mates in isolation, and given that each signaller in a crowd has the potential to flexibly adjust many facets of their signalling bouts, we may expect that signallers will adjust their displays in response to the displays of rivals. For example, if potential mates prefer signallers displaying at high rates and one signaller in a group increases the rate at which they are signalling, this changes the relative attractiveness of every other signaller in the area. Formica et al. (2011) and Oh and Badyaev (2010) both demonstrate that males may move to different social groups in response to the relative attractiveness of rivals, but we may also expect these signallers to adjust their displays in response to changes in their relative attractiveness.

Indeed, evidence suggests that males do change their sexual signals in response to a change in the presence of competition. For example, in the fiddler crab *Austruca annulipes*, males increase the rate at which they wave their major claw as the number of waving neighbours nearby increases above two (Milner et al., 2012). Three-spined sticklebacks (*Gasterosteus aculeatus*) perform more courtship fanning and glueing to females after being regularly presented with a dummy rival male (Kim & Velando, 2014). Male *Drosophila prolongata* in the presence of rival males will switch their courtship from ‘leg vibration’ to ‘rubbing’, which is less conspicuous and therefore less likely to be interrupted by rivals (Setoguchi et al., 2015). Male field crickets (*Gryllus campestris*) increase the performance of their own calling song in response to other males calling, though the opposite is true when callers are present within one metre (Wilde et al., *In press*). This research (as well as Formica et al., 2011; Oh and Badyaev, 2010; Salmon and Atsaides, 1968) investigates how signallers react to the presence or absence of rivals, yet the behaviours of rival signallers may also be important. However, as yet there has been little research investigating whether signallers adjust in response to changes in rival signalling behaviour, yet this is an important step in understanding the extent of the flexibility of these displays, as well their the proximate and ultimate explanations.

Here we investigate how male fiddler crabs (*Afruca tangeri*) adjust their bout-structured sexual signals in response to fine-scale changes in the display behaviour of a biomimetic robotic rival male. Male *A. tangeri* have an enlarged claw (major claw) that they wave repeatedly to attract females. Their waving displays are clearly bout-structured, as waves (raising and lowering of the claw, lasting one second or less) are interspersed with pauses that may be very brief (sometimes less than one second, within signalling bout) or much longer (between signalling bouts). Males may also alternate between periods of high-intensity waving (high speed, short inter-wave interval, high claw elevation) and low-intensity waving (slower speed, longer inter-wave interval, lower claw elevation). This makes *A. tangeri* ideal for investigating adjustments to bout-structured displays.

We hypothesise that focal males will adjust their signalling behaviour in response to changes in the signalling behaviour of the biomimetic rival. Specifically, we predict that males will court at a faster rate and for longer bouts when the robotic rival is courting, as this strategy matching may be a means of remaining competitive in the estimations of potential mates. This may also occur as males may eavesdrop on rival male signalling to infer the location of mate-searching females, as has been shown in wolf spiders (*Schizocosa ocreata*: Clark et al., 2012) and guppies (*Poecilia reticulata*: Webster and Laland, 2013).

## 2 Methods

Using a robot made to mimic the signalling behaviour of male *A. tangeri*, we independently manipulated static (claw size) and dynamic (wave speed) aspects of a perceived rival’s signals and monitored the behaviour of male *A. tangeri* to investigate whether any changes in their displays or time spent in burrow were related to changes in the robotic rival’s display behaviour.

### 2.1 Study area and species

The study was carried out on a wild population of *A. tangeri* that inhabit the Ria Formosa Natural Park on the south-eastern coast of Portugal between May and July 2022. At low tide, this area consists mainly of mudflats, salt marshes and dry sandy banks. The data for this study were collected between Manta Rota and Cacela Velha (37.16 N, 7.53W).

*A. tangeri* live on coastal mudflats and sandbanks and during low tide in the mating season they emerge from their subterranean burrows to maintain their burrow, forage and either perform sexual signals (males) or search for and evaluate potential mates (females). Females have been shown to prefer males with a larger claw (Oliveira & Custódio, 1998) and, in other species, a faster waving rate (Callander et al., 2012). When a female closely inspects a male who is courting, he will run to his burrow entrance and drum his major claw on his carapace and often enter his burrow and drum below ground. The vibrations produced when drumming below ground are thought to provide information to the female on his burrow characteristics (Mowles et al., 2017). If the female wishes to inspect the male’s burrow, she will enter, after which she will either copulate with the male and remain in his burrow to lay her clutch of fertilised eggs, or leave. If she chooses to mate and stay in the burrow, the male will leave and steal or dig another burrow (Crane, 1975; Wolfrath, 1993).

In addition to using their burrows for attracting and mating with females, *A. tangeri* males use their burrows as refuges to wet their gills, escape from predators and avoid aggression from neighbours (Crane, 1975). While in their burrow, males cannot see any mate-searching female or wave. Therefore, we consider the time a male spends in his burrow as an aspect of his displaying behaviour when he enters to solicit burrow inspection by nearby females (Wolfrath, 1993) or when entering at other times, which may occur as a way to reduce costs when his sexual signals are ineffective or provoke increased aggression from neighbouring rivals.

### 2.2 Robotic stimulus male

A 3D scan of a fiddler crab specimen from a museum collection is available to download and use at threedscans.com/crab/fiddler-crab (Laric, 2016). The body and major claw of these scans were 3D-printed separately by FabLab Devon (fablabdevon.org). The major claws were printed in two sizes: one where the major claw was 5cm from manus base to pollex tip (the approximate mean length of male *A. tangeri* major claws in our records, JAW unpubl. - see supplementary materials), and one where the major claw measured 7cm (the largest major claw length in our records, JAW unpubl.). The body was 3D printed in one size: the size of the original scan when the major claw was scaled to the mean length. After printing, the models were painted that closely resemble male *A. tangeri* colours to the human eye using acrylic paint. These models can be seen in figure 1.

**Figure 1:**
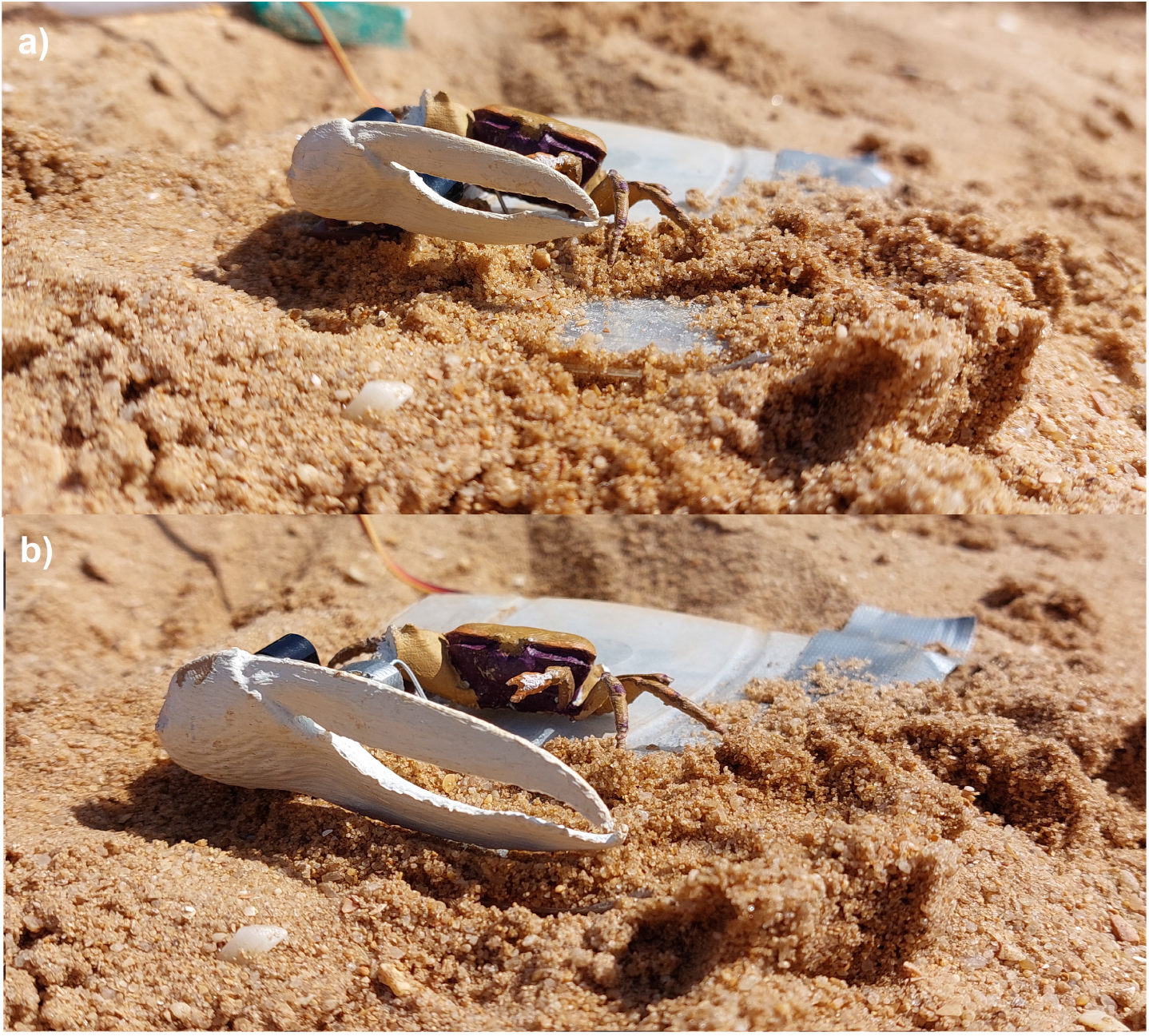
Biomimetic robotic rival male *Afruca tangeri* with the small claw attached (a) and the large claw attached (b).

The printed major claw was glued to a mobile joint that allowed vertical movement. This joint was then fixed to a plastic base and wire passed through the plastic base, allowing the printed claw to be joined to a Tower Pro Micro Servo SG90. This micro servo was attached to an Arduino Uno Rev3 microcontroller which controlled the timing of the micro servo movement and therefore the vertical movement of the printed claw, emulating the waving of a male *A. tangeri*. The Arduino microcontroller was also connected to an HC-05 Bluetooth serial transceiver, allowing the wave speed of the robot to be changed remotely via Bluetooth connection to a mobile phone (see video in supplementary materials). The robotic stimulus male could wave at two speeds: a “Slow” speed of one wave every two seconds, simulating low-intensity waving; and a “Fast” speed of one wave every one second, simulating high-intensity waving. These wave speeds were estimated using video data collected from previous field seasons (JAW unpubl.).

We observed multiple male *A. tangeri* aggressively challenging the robotic rival during test trials. Given that we have only observed *A. tangeri* fighting with other conspecifics, the focal males perceived the robotic rival as a conspecific rival male.

### 2.3 Experimental set-up

All experimental trials were carried out during diurnal low tide periods. Before the trial began, a male who was actively waving on the surface next to his burrow was selected as the focal male. All burrows within a 60cm radius of the focal’s burrow were plugged using bundles of samphire. This prevents any crabs that occupy these burrows from emerging onto the surface for a short period of time and temporarily removes all but the focal male from the experimental area (all burrows were unplugged after the trial was complete). The robotic stimulus male was then placed on the substrate 30cm away from the focal male’s burrow entrance (which is approximately the average distance to the nearest neighbouring burrow, JAW unpubl.) with either the “small” or “large” claw attached. Each focal male was only presented with one claw size for the duration of their trial. The presentation of robot claw size was randomised across males.

Two GoPro Hero 4 cameras were then set up to record the behaviour of the focal male. One GoPro was placed behind the robotic stimulus and the other placed on the other side of the focal male’s burrow, facing the robotic stimulus. This camera set up ensured that the frontal view of the focal male was visible at all times.

Once all equipment was set up, the experimenter (JAW) retreated to a distance at which his presence would not affect focal male behaviour (*>*10 metres) and a timer was started. If the focal male did not emerge from his burrow within 7.5 minutes, that particular trial was abandoned. If the focal male did emerge from his burrow before this time threshold, the timer was reset and a 5-minute timer was started. This marked the beginning of the first condition of the trial where the robotic stimulus was present but not waving (“No wave” treatment). After 5 minutes of this condition, the robotic stimulus was started waving at one of the two wave speeds (either “Slow wave” or “Fast wave” treatment). After 5 minutes of this condition, the robot waving was stopped and the focal male experienced another “No wave” condition lasting 5 minutes. The final condition was 5 minutes of the robot wave speed that the focal male had not yet experienced. All changes to robot wave speed were done remotely to avoid disturbing the focal male. The time at which the trial started was noted, as well as the substrate surface temperature (measured using Magnusson KC-180A1 infrared thermometer, *±*2°C). The claw size treatment and order of wave speed presentation were balanced across focal males using a randomised-blocks design: trials were conducted in blocks of four (large, Slow-Fast; small, Slow-Fast; large, Fast-Slow; small, Fast-Slow), with the order of these trials fully randomised within each block. The same “small” and “large” claws were used across all males in each claw size treatment and the same carapace was presented to all focal males.

### 2.4 Focal male morphometrics

Once a focal male’s trial had ended, the robotic stimulus, both GoPros and any burrow plugs were removed from the area and a tube trap (Wolfrath, 1993) was placed in the entrance to the focal male’s burrow to capture the focal male. If the focal male entered the trap, we measured the distance between the base of his manus and the tip of his pollex (claw length) and the widest point of the dorsal side of his carapace (carapace width) using a Whitworth electronic caliper (*±* 0.01mm), and weighed him using Ohaus Traveller’s scales (*±*0.1g).

If the focal male did not enter the trap within 20 minutes (n=38), the trap was removed and his claw length and carapace size were estimated from the GoPro footage using the software ImageJ (Schneider et al., 2012), with a measuring tape placed above his burrow for scale.

The experimental trials were carried out on 70 focal males, but only 58 were used for the final analysis as some trials were either interrupted by human passers-by or the focal males left the experimental area during the trial. Table 1 shows the focal male and trial information for each of the 4 possible experimental treatments.

**Table 1:** Information about the trials in each of the 4 possible experimental conditions. Claw length and carapace width refer to the focal male.

| Robot claw size | Presentation order | Number of trials | Mean claw length | Claw length SD | Mean carapace width | Carapce width SD | Mean temperature | Temperature SD |
| --- | --- | --- | --- | --- | --- | --- | --- | --- |
| Large claw | Fast/Slow | 14 | 44.4 | 11.2 | 25.9 | 4.1 | 24.3 | 3.6 |
| Large claw | Slow/Fast | 13 | 45.7 | 9.5 | 26.8 | 4.8 | 24.6 | 4.2 |
| Small claw | Fast/Slow | 18 | 41.9 | 12.5 | 26.4 | 4.8 | 23.7 | 3.4 |
| Small claw | Slow/Fast | 14 | 43.6 | 10.6 | 27.2 | 4.2 | 24.3 | 3.3 |

### 2.5 Ethical note

This study uses biologically realistic interventions in a natural environment. Captured individuals were returned to their burrow after measurement and neighbouring burrows were unplugged after the trial had ended. All work was carried out with approval from the University of Exeter’s Research Ethics Panel (application ID: 513844) and the Instituto da Conservacão da Natureza e das Florestas (permit ID: 579 / 2022 / CAPT).

### 2.6 Behaviour coding and data

For each experimental trial we extracted all time points at which a male entered or exited his burrow, performed a wave with his major claw, performed a threat display or fought a rival male, and when a female was visible in either of the two videos (the definitions for these behaviours are in table 2). We then calculated the proportion of time a focal male spent in his burrow during each wave speed treatment.

**Table 2:** Ethogram showing detailed definitions of the behaviours coded and used in subsequent analyses.

| Behaviour | Type | Description |
| --- | --- | --- |
| Aggressive stance | Point event | Male opens and raises claw in aggressive stance. (Not waving) |
| Wave | Point event | Male waving enlarged claw above the level of their eye and returns claw to resting position. |
| In burrow | State event | All of males legs are not visible in camera footage. |
| Fight | State event | Male makes physical contact with another individual. |
| Female is visible in camera footage. | Point event | Female enters frame. |
| Female exits camera footage | Point event | Female leaves frame. |

Videos were coded once by one of the research team (JAW) who was not blind to the claw size or wave speed treatment. Ten percent of the trial videos were also coded independently by an individual who was given videos where the robotic rival was not visible and was therefore blind to both the claw size and wave speed treatments. The inter-rater reliability was calculated as 89%.

In their study advocating the use of HMMs when analysing bout-structured courtship displays such as this, Perry et al. (2019) used the continuous time interval between courtship displays as their response variable, modelling this with gamma distributions. For our fiddler crab data, the distributions of time intervals between waves are not identifiable using gamma distributions. Instead, we opted to model the number of waves per five seconds (5 s) using the Poisson distribution. While this does lose some fine-scale information about the precise timing of each wave, we can still use it to estimate the courtship intensity and the latent state of the focal males every 5 s while controlling for temporal autocorrelation between successive intervals.

Therefore, the data were then divided into 5-second time intervals for the purpose of statistical analysis. Any 5-s intervals in which the focal male left the field of view of both video cameras for any portion of the 5 s was discounted. In each of the remaining intervals, we recorded the number of times the focal male waved, the number of females in frame, and whether the focal male aggressed or fought a rival male during those 5 s. We had a total of 10,400 5-s intervals across 58 individuals.

### 2.7 Statistical analyses

#### 2.7.1 Courtship behaviour

The courtship behaviour of *A. tangeri* is bout-structured: a male repeats waves over time before breaking for a period. Therefore, we used Hidden Markov Models (HMMs) as this is a data driven approach that respects the bout structure of the behavioural data (McClintock et al., 2020; Perry et al., 2019) and allows us to model the latent state of focal males.

HMMs are statistical models that assume an individual is in one of a number of possible states at any given time. The state an individual is in is not observable (hidden), but the expression of observable behaviour at any given time is an indicator of the hidden state they are in at that time. In this study, we model focal males as being in one of two hidden states: a signalling state (in which a male is actively courting) and a non-signalling state (in which a male is doing anything other than actively courting but is on the substrate surface). We use the number of waves per 5 s as a proxy from which to infer the underlying state of the male. These models use time-series data to estimate the probabilities that an individual transitions between states (from signalling to non-signalling, or vice versa) from one 5-s time interval to the next. These state transition probabilities can be used to calculate the average length of signalling and non-signalling bouts (average length of bout 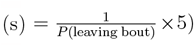). HMMs simultaneously estimate the rate of waving (per 5 s) in each state, which we use as a measure of the intensity of signalling (this primarily applies to the signalling state, which is the state with the higher waving rate by definition). The probability of continuing a signalling bout (or transitioning to a non-signalling bout) and the rate of waving estimated by the HMMs allow us to disentangle multiple ways that a male can adjust his bout-structured waving behaviour, and may reveal flexible changes in signalling that would be overlooked by coarser analytical techniques. The HMMs also allow us to estimate how the state transition probabilities and waving rates depend on predictor variables characterising the male and his social context. We estimate the effect each predictor variable has on (1) the probability a focal male stays in the signalling state, (2) the probability he stays in the non-signalling state, (3) the rate of waving when in the signalling state and (4) the rate of waving when in the non-signalling state. The predictors we included were: the wave speed of the robot, the claw size of the robot, the number of females present in that 5-s interval, whether a rival male was present in that 5-s interval, the temperature at the start of the trial, the focal male’s carapace width, his claw-length-to-carapace-width ratio, and the time of 5-s interval relative to low tide. Random intercepts were included for focal male identity to account for non-independence among measurements and capture natural variation among males in their average transition probabilities and waving rates. A full outline of the linear models used can be found in equations 1 and 2 in the supplementary materials.

See figure 2 as an example of how HMMs predict the latent state of focal individuals.

**Figure 2:**
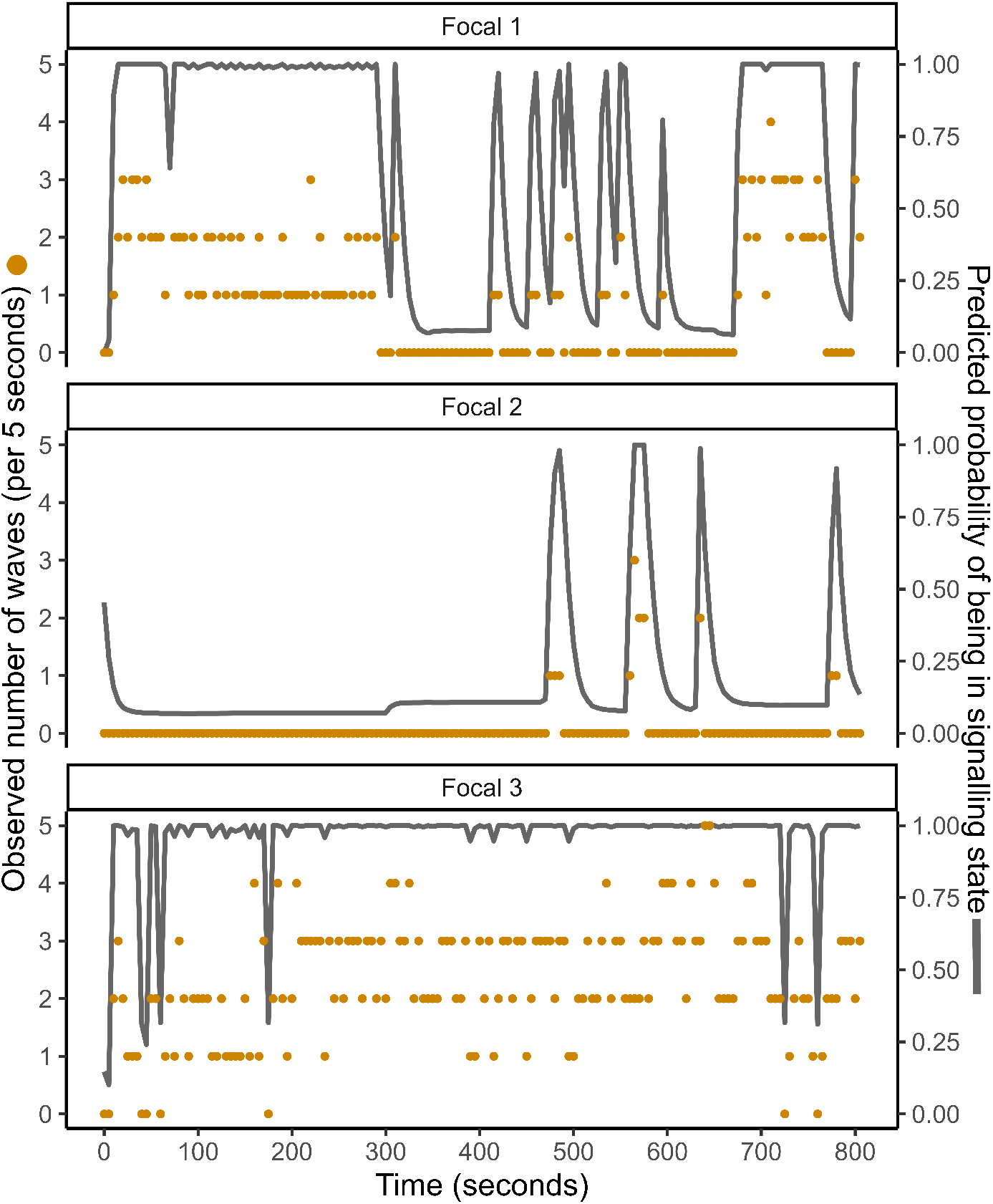
The observed number of waves a focal male performed over 13.5 minutes from 3 example trials and the predicted probability that male is in signalling state based on his behaviour, from the hidden Markov model. The orange points show the number of waves a male performed per 5 seconds (observed data, left y axis) and the grey line shows the predicted probability of being in signalling state (model prediction, right y axis).

#### 2.7.2 Time spent in burrow

We used a linear hurdle model to analyse whether males adjusted the time spent in their burrow in response to changes in the behaviour and claw size of the robotic rival. In this analysis, we asked how the signalling behaviour of the biomimetic rival male affected (1) the probability that a male entered his burrow and (2) how long he spent there. For each of the five-minute treatment conditions (“No wave”, “Slow wave” or “Fast wave”), the probability that the male entered his burrow at all during that five-minute period was modelled using the Bernoulli distribution (binary outcome, the ‘hurdle’). For those cases where the male did enter his burrow, the proportion of the five minutes he spent there was modelled using the Beta distribution (ranging between 0 and 1). The predictors used in both parts of the linear hurdle model were robot claw size, robot wave speed, the focal male’s carapace width, his claw-length-to-carapace-width ratio, the temperature at the start of the trial, the time the trial started relative to low tide, and the treatment order. Random intercepts were included for focal male identity to account for non-independence among measurements and capture natural variation among males in their tendency to enter and spend time in the burrow.

All models were written in Stan (Carpenter et al., 2017) and compiled and run using CmdStanR version 0.5.3 (Gabry & Cešnovar, 2022) and CmdStan version 2.30.1 (Stan Development Team, 2018) in R version 4.1.2 (R Core Team, 2022; RStudio Team, 2020). For more information on HMMs see Glennie et al. (2023), McClintock et al. (2020), and Perry et al. (2019).

The summary of all model outputs, including prior distributions, posterior means, standard errors, and effective sample sizes for all parameters can be found in the supplementary material. The potential scale reduction factor, 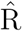 was *≈*1 for all parameters (Gelman & Rubin, 1992). The 89% highest density interval is reported throughout (Kruschke, 2014; McElreath, 2020). All continuous predictors (temperature, time, carapace width, claw:carapace ratio) were converted to z-scores to facilitate model convergence and aid interpretation.

## 3 Results

### 3.1 Hidden Markov Models of signalling behaviour

#### 3.1.1 Wave rate in each state

There is a clearly identifiable difference in wave rate between the two latent states: 0.02 waves per 5-s in the non-signalling state (intercept term posterior mean, 89% HDI [0.01, 0.03]) and 1.4 waves per 5-s in the signalling state (intercept term posterior mean, 89% HDI [1.04, 1.86], see figure 3). The model could therefore identify and predict the hidden state of individuals based on their observable signalling behaviours.

**Figure 3:**
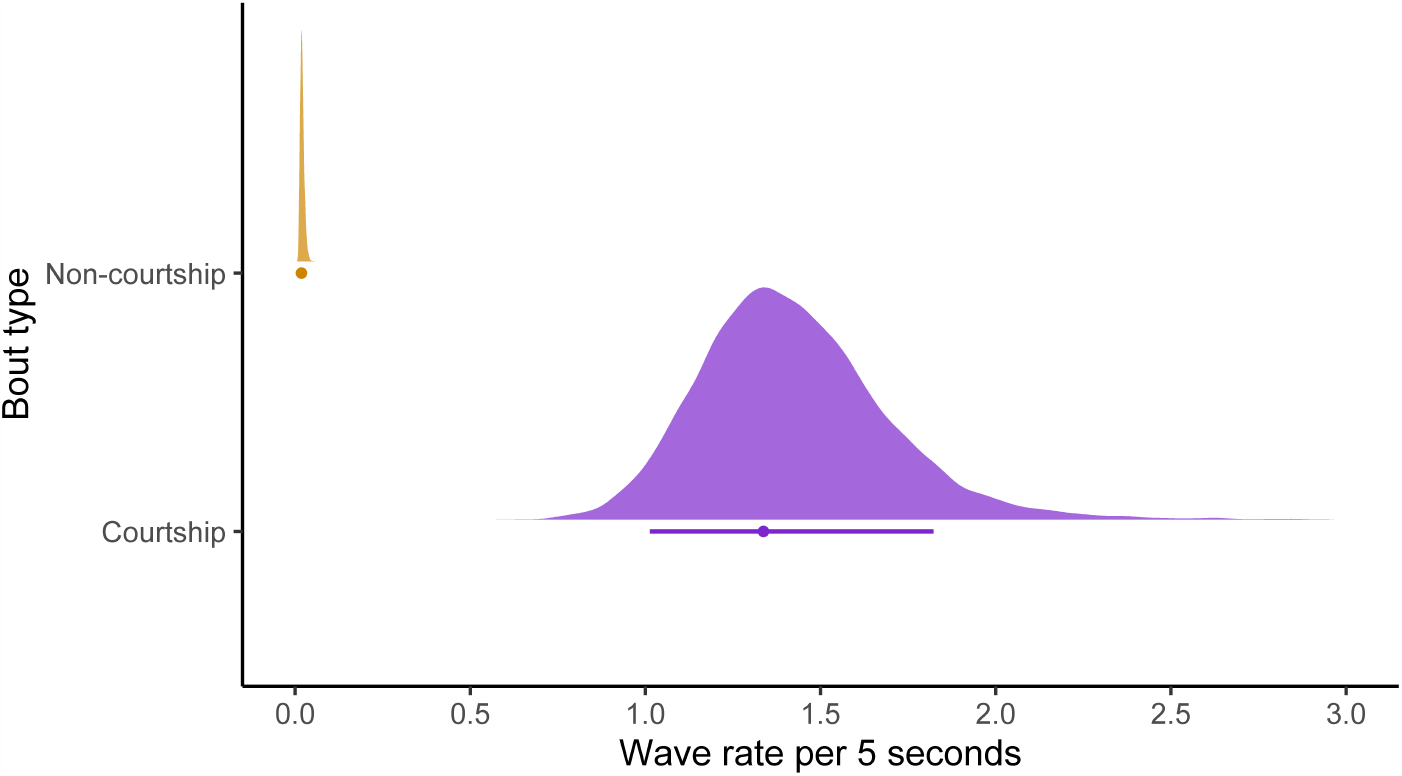
Posterior distributions for the wave rates in both signalling (purple) and non-signalling (orange) state. Below each distribution is the mode (point) and 89% highest density interval (line) of the distribution. These are the baseline rates (intercept) without the effect of any predictors, i.e. for a male of average carapace width and claw size at the average temperature and lowest tide point, in the presence of a non-waving, small-clawed robotic stimulus and no females or rival males.

**Figure 4:**
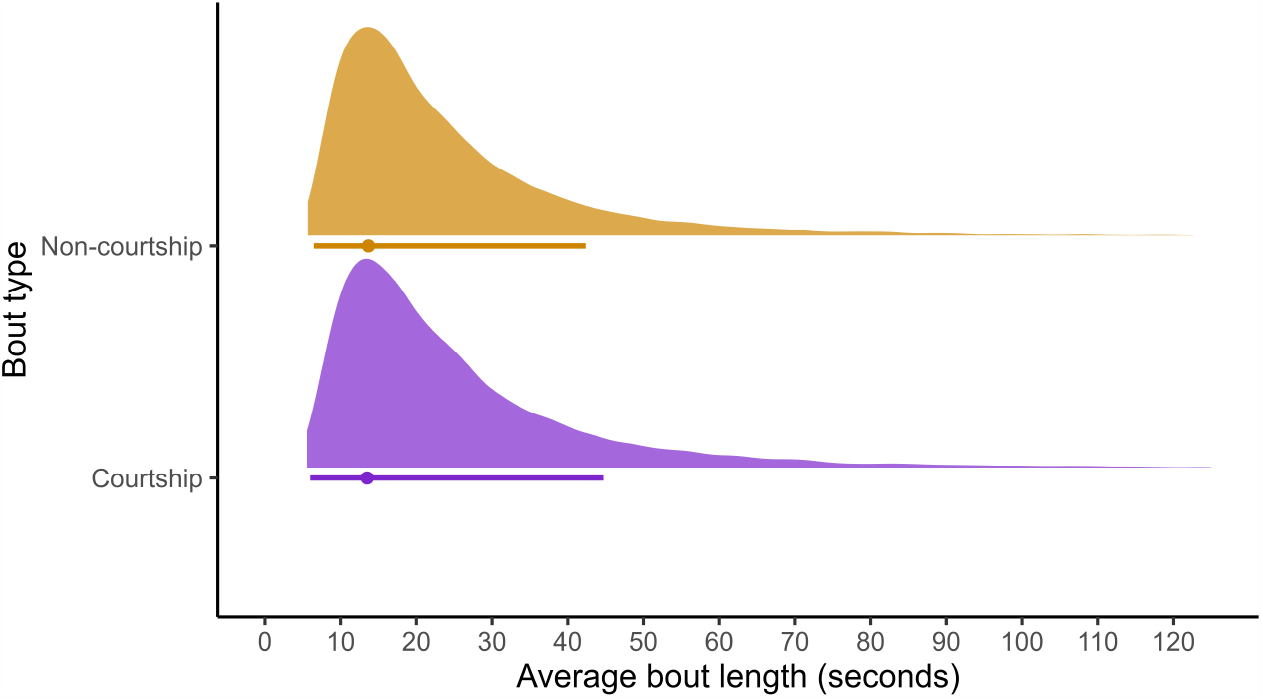
Posterior predictions of average bout length for non-signalling (orange) and signalling (purple). Shown beneath each density plot are the posterior mode (point), and 89% interval (line).

#### 3.1.2 Wave rate in signalling state

There is little evidence to suggest that the robot’s behaviour and claw size or any other predictors had an effect on the rate of waving in the signalling state (see model output tables in supplementary materials). Because the wave rate during non-signalling is close to zero and we had no prior expectation that any of the predictors would affect this, we do not provide information on their estimated effects.

#### 3.1.3 Duration of signalling bouts

Males had a lower probability of transitioning from signalling to non-signalling when the robotic rival stimulus was waving at the “Fast” speed (89% highest density interval (HDI) [-0.02, 2.03]), leading to longer signalling bouts in this treatment condition (see figure 5). There was no apparent effect of the robot waving at the “Slow” speed (89% HDI [-1.09, 1.04]) or the claw size of robot (89% HDI [-1.25, 1.27]).

**Figure 5:**
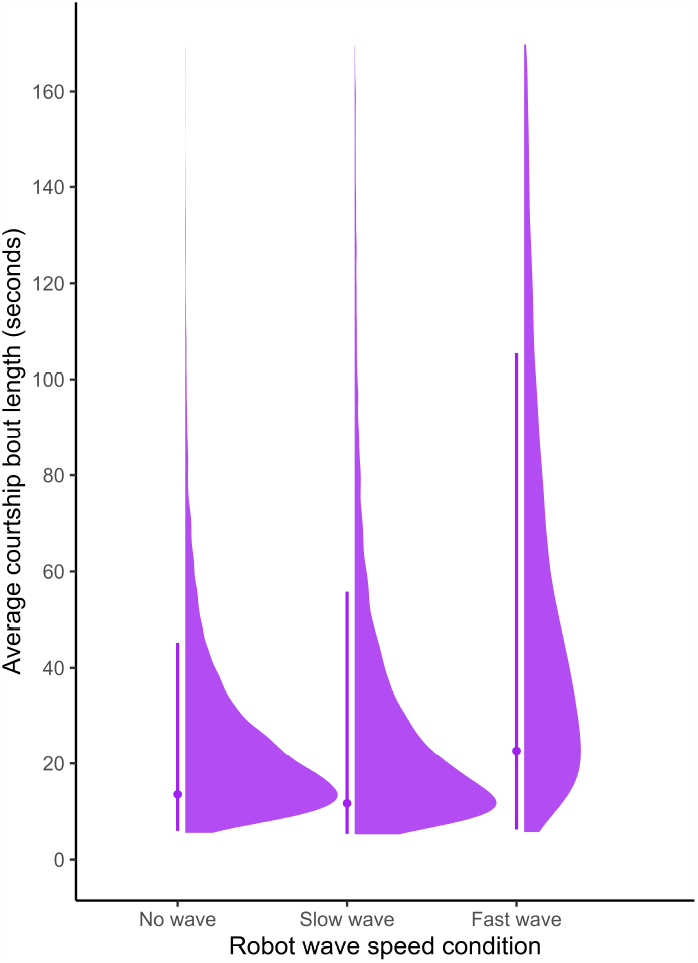
Posterior predictions of average length of signalling bouts in seconds between the three robot wave speed conditions. The density plots show the predicted average signalling bout length under the three wave speed conditions. The points show the posterior mode and the lines span the 89% highest density interval for each wave speed condition.

There is little evidence to suggest any other predictors had an effect on the probability of remaining in a signalling bout (see model output tables in supplementary materials).

#### 3.1.4 Duration of non-signalling bouts

The behaviour and claw size of the robotic stimulus did not appear to affect the probability of remaining in a non-signalling bout (89% HDI “Slow wave” [-1.32, 0.81], “Fast wave” [-1.49, 1.19], “Large claw” [-1.29, 1.28]) and non-signalling bouts therefore lasted similar times across all robot treatments. There is also little evidence to suggest that any other predictors affected the probability of remaining in a non-signalling state (see model output tables in supplementary materials).

### 3.2 Linear hurdle model of entering and spending time in burrow

#### 3.2.1 Probability of entering the burrow

Males were less likely to enter their burrow when the robot was waving, but the difference was the same across both wave speeds: “Slow” speed (89% HDI [-3.52, -2.06]) and “Fast” speed (89% HDI [-3.58, -2.09], see open vs. closed circles in figure 6). Therefore, there was an effect of the robot waving vs. not, but not of the wave speed. The claw size of the robot had no effect on the probability a male entered his burrow (89% HDI [-0.78, 0.56]). No other predictors included showed any effect (see model output tables in supplementary materials).

**Figure 6:**
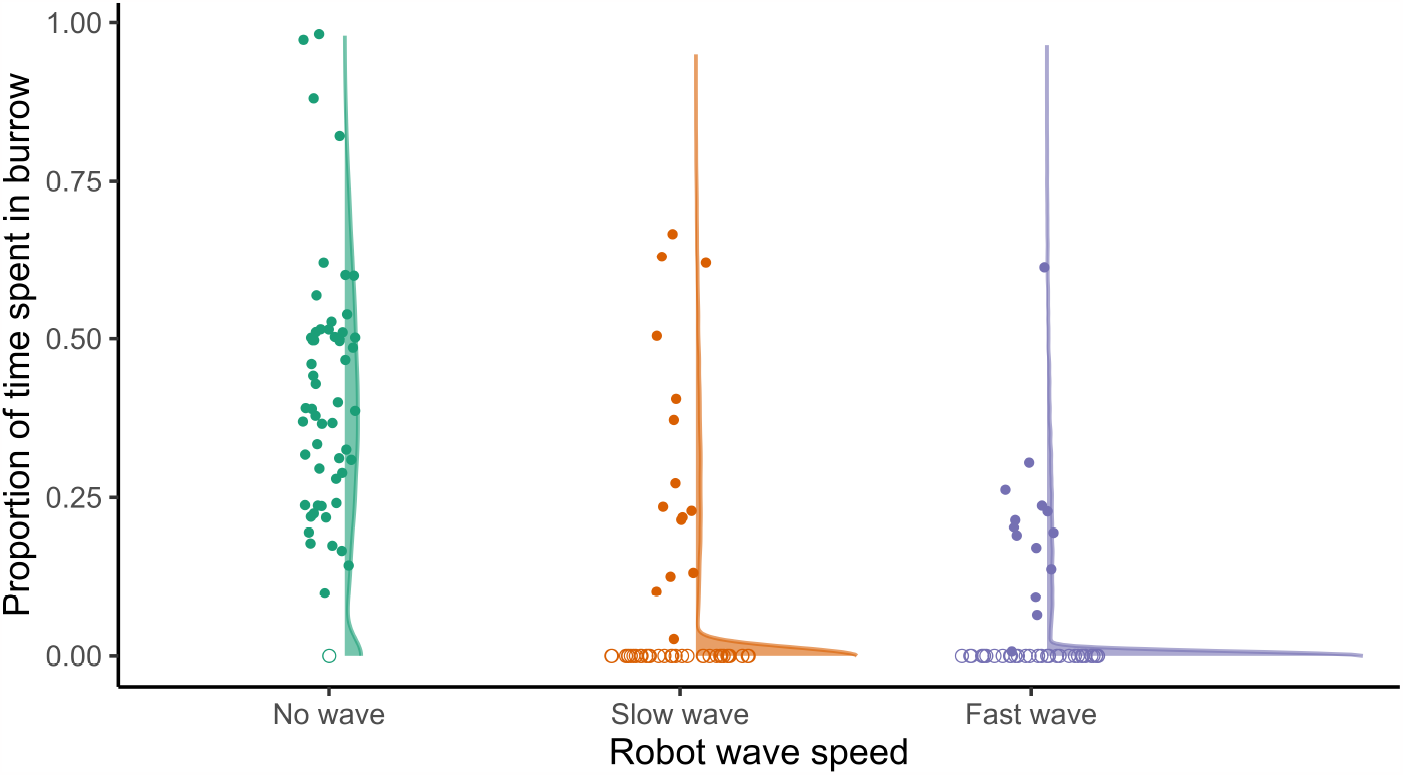
Raw data (circles) and model predictions (density plots) of the proportion of time males spent in their burrow during each wave speed treatment. The distributions on the right show the model predictions. The points on the left show the proportion of time males spent in their burrow (filled circles), with open circles if they did not enter their burrow at all during that treatment (from the raw data).

#### 3.2.2 Proportion of time spent in burrow

Males spent less time in their burrow when the robot was waving, either “Slow” (89% HDI [-0.77, -0.01]) or “Fast” (89% HDI [-1.20, -0.37]) and there is evidence to suggest that males spent the least time in their burrows during the “Fast” wave speed condition (see figure 6). When the robot had a large claw, males spent more time in their burrow than when the robot had a small claw (89% HDI [0.03, 0.74]).

Males also spent less time in their burrow at warmer temperatures (89% HDI [-0.39, -0.04]) and there is evidence to suggest that males spent less time in their burrows when the “Fast” wave speed condition was presented first (89% HDI [-0.71, -0.003]). No other predictors had a clear effect (see model output tables in supplementary materials).

## 4 Discussion

Studies investigating the tactical adjustment of signalling behaviours have, thus far, mainly focused on how signallers adjust their signalling in response to the varying presence of rivals (Kim & Velando, 2014; Milner et al., 2012; Setoguchi et al., 2015; Wilde et al., *In press*). In this experimental study of wild fiddler crabs, we investigated how males adjust their signalling behaviour in response to fine-scale changes in both static (claw size) and dynamic (wave rate) aspects of rival displays. We used an analytical approach that provides a holistic, detailed view of how males adjust multiple aspects of their bout-structured sexual signals and how they invest in time spent in visual contact with conspecifics. We found that males extended their display bouts when a robotic rival was waving at a fast rate, but found no evidence to suggest that males adjusted their wave rate within display bouts in response to changes in rival behaviour. We also found that males were less likely to enter their burrow when a rival was waving at high intensity and that, if they did enter their burrow, they spent less time in there if the rival was waving. Males also spent less time in their burrow if the rival had a small claw.

Males were less likely to end a signalling bout when the robot was waving fast, meaning the signalling bouts lasted longer during this treatment. Given that male *A. tangeri* usually only perform the “Fast” wave when females are close (How et al., 2008), it seems likely that the focal males were eavesdropping on the wave rate of the robotic rival and using this as a cue to nearby females. Males eavesdropping on the signalling of rival males to gain information about females has been documented in guppies (*P. reticulata*; Webster and Laland, 2013), wolf spiders (*Schizocosa ocreata*; Clark et al., 2012), whitethroats (*Sylvia communis*; Balsby and Dabelsteen, 2005), and an Australian species of fiddler crab (*Austruca mjoebergi* ; Milner et al., 2010). By using the behaviour of rival conspecifics to effectively extend the distance at which mate-searching females can be detected, males can tactically adjust their signalling behaviour to increase effort in signalling only when it is most effective in attracting females. We also found that males were less likely to enter their burrow when the robot was waving at any speed, and spend less time underground if they did enter, compared to when the robot was not waving. Again it seems likely that this is due to rival signalling potentially indicating female proximity, as it is in the best interests of a male not to enter his burrow when there is an immediate opportunity to attract a mate. The finding that experimental treatment order affected the amount of time males spent in their burrows, with males spending less time in their burrow overall when presented with the “Fast” wave speed condition first may also fit into this conclusion. If males are using the fast waving of the robotic as an indicator of females nearby, males may strategically spend less time in their burrow for some time after the “Fast” wave speed condition so as to avoid missing any mate-searching females that may be wandering through the area.

Given that there was no detectable effect of the wave speed treatment or the robot’s claw size on the focal wave rate, we did not find evidence that *A. tangeri* tactically adjust their waving intensity in response to changes in their relative attractiveness caused by changes in rival signalling. This finding fits with previous work showing that male *A. tangeri* do not change their wave rate in response to video playback of a male waving at mostly slow speeds (Burford et al., 2000), and that male *A. annulipes* do not change their wave rate in response to 1-2 waving rivals (Milner et al., 2012). Given that changes in rival signalling behaviour will typically alter the efficacy of a male’s own displays, we might expect him to adjust his signalling behaviour accordingly, however this may only be true with larger number of rivals. Formal models are needed to identify evolutionarily stable strategies for signalling in such competitive situations.

Males spent a larger proportion of the trial period in their burrow when the robot had a large claw. This may be a strategy to avoid aggression from highly competitive conspecifics. Fiddler crabs use their burrows as refuges from conspecific aggression and when an invading male is attempting to steal a burrow, the burrow owner will often enter the burrow to avoid attack and to prevent his burrow being stolen (Muramatsu & Koga, 2016). There is some evidence to suggest that males use rival claw size as a means of assessing fighting ability (Lailvaux et al., 2009, but see Morrell et al., 2005). Therefore, if males perceive the robot as a threat, they may spend longer in their burrow as a defensive strategy. Males may also spend longer in their burrows when the robot has a large claw as they may be less likely to out-compete large-clawed males in attracting a female, and so may strategically spend time sheltering in their burrow as a result. This finding suggests that males do adjust some aspects of their behaviour in response to the static weapon/signalling ornament of rivals, though not necessarily the timing of their bout-structured signalling. By analysing an individual’s signalling strategy *in situ*, as we have done here, we can begin to disentangle some of the complex and sometimes antagonistic effects of the signaller’s social environment.

The use of HMMs allowed us to infer the underlying state of the focal male (signalling vs. non-signalling) and any adjustments he made to the bout structure of his displays. These models worked well in estimating state (see figure 2) based on the sequence of wave rates per 5-second period for each individual, and identified the wave rate within each state well (see figure 3). These models are being used more frequently in ecology (Glennie et al., 2023; McClintock et al., 2020). They are a powerful tool for analysing the underlying state dynamics of many systems and future research should consider them as viable analytical options.

While a powerful tool, HMMs do require a sizeable amount of clean data to pick up on more subtle effects or the effects of events that are sparsely represented in the data. For example, we found no strong evidence that male signalling was affected by females wandering through the trial area. Studies on other fiddler crab species have shown that males do react to females in the wild by waving at a higher intensity (Burford et al., 2000; Crane, 1975; Milner et al., 2010), so we expected to see an effect of female presence in our data. However, females were present in the trial area for only 4.7% of the 5-s periods. It is possible that we would have been able to detect an effect of females if they had been better represented in the sample, but this was not the focus of our experimental study. Nonetheless, we were able to control for any effect of female presence while investigating the effects of the robotic rival on signalling state and structure.

In summary, our study shows that male fiddler crabs adjust the length of their signalling bouts in response to high-intensity waving from a nearby rival and this is likely due to rival signalling being used as a cue to nearby females. However, we found no evidence that males tactically adjust their signalling intensity in response to changes in their relative attractiveness caused by rival behaviour. In light of these findings, our study highlights the complex interplay between male fiddler crabs’ sexual signals and their response to both static and dynamic characteristics of rival males. By examining the flexibility of male *A. tangeri* signalling behaviour in response to a biomimetic rival, we contribute valuable insights into the strategies that shape sexual signalling in a competitive environment. As our understanding of such flexible and dynamic signalling displays continues to grow, we hope that this work will not only inform future research on the proximate and ultimate explanations of signalling behaviour in fiddler crabs, but also inspire broader investigations into the role of competition and flexibility in shaping animal communication systems across diverse taxa. Future research should integrate these findings into a broader theoretical evolutionary framework to understand what forms of tactical signalling adjustment we would expect to evolve, and under what conditions.

## 5 Acknowledgments

Thanks to the Instituto da Conservação da Natureza e das Florestas for allowing this work to be done. Many thanks to Beki Hooper, Lorna Wilde and ChatGPT for advice and feedback on this manuscript, as well as the rest of the CRAB group for helpful discussions. Massive thanks to our crab-artist-in-residence Beki Hooper for her work painting the 3D models. This work was supported by the Natural Environment Research Council via a GW4+ DTP2 studentship (NE/S007504/1). Thank you to Adam Chee for independent video coding.

## Notes

### Competing Interest Statement

The authors have declared no competing interest.

