## Supplementary materials for "Biomimetic robots reveal flexible adjustment of sexual signalling in a wild invertebrate"

### 6 Supplementary materials

Video of biomimetic robot rival - https://youtu.be/K3CI4HynAFY

$$
\begin{aligned}
Pr(s_t = i \mid s_{t-1} = i) &\sim \text{Bernoulli}(p_i) \\
\text{logit}(p_i) &= \alpha_{i,z} + \beta_{1,i}x_1 + \beta_{1,i}x_2 + ... + \beta_{n,i}x_n \\
\alpha_{i,z} &\sim \text{Normal}(\mu_i, \sigma) \\
\mu_i &\sim \text{Normal}(0, 1) \\
\sigma &\sim \text{Exponential}(2) \\
\beta_{n,i} &\sim \text{Normal}(0, 0.8)
\end{aligned} \tag{1}
$$

$$
\begin{aligned}
w_s &\sim \text{Poisson}(\lambda_s) \\
\log(\lambda_s) &= \alpha_{s,z} + \beta_{1,s}x_1 + \beta_{1,s}x_2 + ... + \beta_{n,s}x_n \\
\alpha_{s,z} &\sim \text{Normal}(\mu_s, \sigma) \\
\mu_{s\,=\,\text{courtship}} &\sim \text{Normal}(0.5, 0.5) \\
\mu_{s\,=\,\text{non}-\text{courtship}} &\sim \text{Normal}(-4, 0.3) \\
\sigma &\sim \text{Exponential}(10) \\
\beta_{n,s} &\sim \text{Normal}(0, 0.1)
\end{aligned} \tag{2}
$$

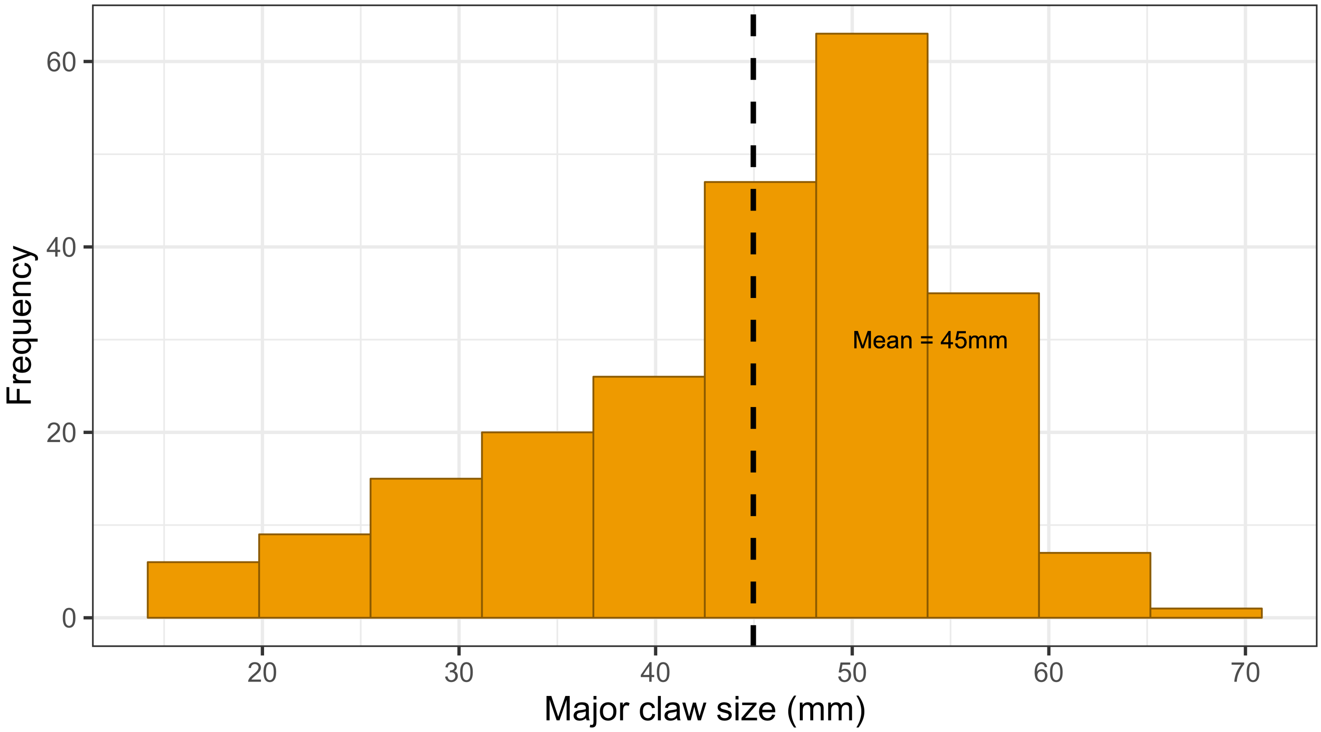


Figure 7: The distribution of 229 male *Afruca tangeri* major claw sizes in millimeters, measured between 2014 - 2022. The dashed line indicates the mean value of 45 mm.

| | Posterior values | | | 89% Highest Density Limits | | | Effective sample size | |
|---|---|---|---|---|---|---|---|---|
| Parameter | Mean | Median | SD | Lower | Upper | R-hat | Bulk | Tail |
| **Wave rates** | | | | | | | | |
| **Intercept mean** | | | | | | | | |
| Signalling state | 1.401 | 1.398 | 1.207 | 1.039 | 1.859 | 1 | 9044 | 5948 |
| Non-signalling state | 0.020 | 0.020 | 1.339 | 0.012 | 0.032 | 1 | 21111 | 6783 |
| **Intercept SD** | | | | | | | | |
| Signalling state | 1.101 | 1.069 | 1.099 | 1.000 | 1.234 | 1 | 11550 | 6398 |
| Non-signalling state | 1.105 | 1.073 | 1.103 | 1.000 | 1.242 | 1 | 13545 | 7060 |
| **Predictors** | | | | | | | | |
| **signalling state** | | | | | | | | |
| Number of females | -0.001 | 0.000 | 0.097 | -0.154 | 0.155 | 1 | 22961 | 8332 |
| Rival presence | -0.040 | -0.041 | 0.098 | -0.191 | 0.122 | 1 | 21987 | 8725 |
| Slow wave speed | 0.009 | 0.009 | 0.092 | -0.146 | 0.149 | 1 | 23352 | 8326 |
| Fast wave speed | 0.046 | 0.047 | 0.087 | -0.087 | 0.189 | 1 | 22232 | 9143 |
| Large claw | 0.000 | 0.002 | 0.100 | -0.164 | 0.158 | 1 | 23650 | 8423 |
| Temperature | 0.003 | 0.002 | 0.097 | -0.146 | 0.165 | 1 | 21951 | 8868 |
| Carapace width | -0.004 | -0.004 | 0.100 | -0.160 | 0.156 | 1 | 20721 | 8384 |
| Claw length:carapace width ratio | -0.004 | -0.006 | 0.099 | -0.159 | 0.156 | 1 | 19500 | 9181 |
| Time relative to low tide | 0.009 | 0.009 | 0.100 | -0.151 | 0.166 | 1 | 24702 | 8018 |
| **Non-signalling state** | | | | | | | | |
| Number of females | -0.001 | -0.001 | 0.100 | -0.167 | 0.154 | 1 | 25213 | 7812 |
| Rival presence | -0.001 | -0.001 | 0.100 | -0.173 | 0.147 | 1 | 23725 | 6116 |
| Slow wave speed | 0.004 | 0.004 | 0.100 | -0.154 | 0.163 | 1 | 20509 | 7185 |
| Fast wave speed | -0.001 | 0.000 | 0.100 | -0.169 | 0.151 | 1 | 23089 | 8064 |
| Large claw | 0.000 | -0.001 | 0.102 | -0.161 | 0.165 | 1 | 21159 | 7020 |
| Temperature | -0.004 | -0.004 | 0.097 | -0.155 | 0.154 | 1 | 24850 | 8333 |
| Carapace width | 0.005 | 0.006 | 0.099 | -0.158 | 0.158 | 1 | 18766 | 7963 |
| Claw length:carapace width ratio | 0.004 | 0.003 | 0.098 | -0.150 | 0.163 | 1 | 21605 | 8016 |
| Time relative to low tide | 0.000 | 0.000 | 0.099 | -0.153 | 0.162 | 1 | 21347 | 8638 |

Table 3: Summary statistics for the Hidden Markov model results looking at how the robotic rival affects the wave rate within each state (courtship/non-courtship). Shows summary statistics of the posterior distributions, $\hat{R}$, and effective sample sizes for the mean of intercepts, standard deviation of intercepts and predictor effect sizes.

| | Posterior values | | | 89% Highest Density Limits | | | Effective sample size | |
|---|---|---|---|---|---|---|---|---|
| **Parameter** | **Mean** | **Median** | **SD** | **Lower** | **Upper** | **R-hat** | **Bulk** | **Tail** |
| **Transition probabilities** | | | | | | | | |
| **Intercept mean** | | | | | | | | |
| Continuing signalling | 0.746 | 0.750 | 0.698 | 0.450 | 0.921 | 1 | 12279 | 5506 |
| Continuing non-signalling | 0.737 | 0.739 | 0.691 | 0.440 | 0.912 | 1 | 14205 | 6868 |
| **Intercept SD** | | | | | | | | |
| Continuing signalling | 0.647 | 0.604 | 0.641 | 0.500 | 0.795 | 1 | 10023 | 7011 |
| Continuing non-signalling | 0.641 | 0.601 | 0.637 | 0.500 | 0.783 | 1 | 9565 | 6666 |
| **Predictors** | | | | | | | | |
| **Continuining signalling** | | | | | | | | |
| Number of females | -0.008 | -0.011 | 0.806 | -1.214 | 1.346 | 1 | 22049 | 7979 |
| Rival presence | -0.043 | -0.048 | 0.818 | -1.395 | 1.239 | 1 | 21412 | 8176 |
| Slow wave speed | 0.028 | 0.017 | 0.668 | -1.091 | 1.044 | 1 | 22984 | 8192 |
| Fast wave speed | 1.013 | 1.005 | 0.644 | -0.018 | 2.032 | 1 | 21150 | 8476 |
| Large claw | -0.003 | 0.004 | 0.794 | -1.245 | 1.274 | 1 | 21649 | 8309 |
| Temperature | -0.425 | -0.426 | 0.761 | -1.611 | 0.790 | 1 | 20071 | 8390 |
| Carapace width | 0.531 | 0.525 | 0.764 | -0.663 | 1.756 | 1 | 20662 | 6868 |
| Claw length:carapace width ratio | 0.419 | 0.416 | 0.778 | -0.803 | 1.701 | 1 | 23844 | 8210 |
| Time relative to low tide | -0.255 | -0.250 | 0.810 | -1.503 | 1.081 | 1 | 20629 | 8370 |
| **Continuining non-signalling** | | | | | | | | |
| Number of females | 0.000 | -0.013 | 0.785 | -1.236 | 1.260 | 1 | 26012 | 8795 |
| Rival presence | 0.061 | 0.068 | 0.780 | -1.139 | 1.351 | 1 | 24628 | 8474 |
| Slow wave speed | -0.251 | -0.265 | 0.670 | -1.321 | 0.813 | 1 | 21813 | 8552 |
| Fast wave speed | -0.246 | -0.255 | 0.831 | -1.488 | 1.185 | 1 | 22215 | 8197 |
| Large claw | -0.001 | -0.005 | 0.801 | -1.291 | 1.284 | 1 | 23490 | 7534 |
| Temperature | -0.422 | -0.424 | 0.768 | -1.622 | 0.815 | 1 | 16635 | 6802 |
| Carapace width | 0.501 | 0.505 | 0.763 | -0.730 | 1.685 | 1 | 16943 | 7780 |
| Claw length:carapace width ratio | 0.390 | 0.398 | 0.773 | -0.842 | 1.644 | 1 | 21890 | 8050 |
| Time relative to low tide | -0.457 | -0.455 | 0.773 | -1.619 | 0.860 | 1 | 22941 | 8204 |

Table 4: Summary statistics for the Hidden Markov model results looking at how the robotic rival affects the probability a male continues in each bout (courtship/non-courtship). Shows summary statistics of the posterior distributions, $\hat{R}$, and effective sample sizes for the mean of intercepts, standard deviation of intercepts and predictor effect sizes.

| | Posterior values | | | 89% Highest Density Limits | | | Effective sample size | |
|---|---|---|---|---|---|---|---|---|
| **Parameter** | **Mean** | **Median** | **SD** | **Lower** | **Upper** | **R-hat** | **Bulk** | **Tail** |
| **Probability of entering burrow** | | | | | | | | |
| Intercepts mean | 0.866 | 0.865 | 0.606 | 0.764 | 0.926 | 1 | 26223 | 18523 |
| Intercepts SD | 0.757 | 0.748 | 0.432 | 0.001 | 1.306 | 1 | 4923 | 8410 |
| **Predictors** | | | | | | | | |
| Slow wave | -2.796 | -2.786 | 0.464 | -3.522 | -2.058 | 1 | 21430 | 18186 |
| Fast wave | -2.841 | -2.831 | 0.470 | -3.581 | -2.091 | 1 | 21781 | 19658 |
| Large claw | -0.099 | -0.101 | 0.422 | -0.781 | 0.560 | 1 | 29801 | 18552 |
| Temperature | 0.532 | 0.531 | 0.559 | 0.439 | 0.624 | 1 | 35921 | 18732 |
| Carapace width | 0.147 | 0.145 | 0.324 | -0.365 | 0.667 | 1 | 26447 | 18484 |
| Claw:carapace ratio | 0.152 | 0.154 | 0.330 | -0.356 | 0.684 | 1 | 24956 | 19090 |
| Time relative to low tide | -0.040 | -0.039 | 0.239 | -0.415 | 0.346 | 1 | 33946 | 18743 |
| Treatment order | 0.403 | 0.395 | 0.432 | -0.295 | 1.071 | 1 | 30587 | 17005 |
| **Proportion of period in burrow** | | | | | | | | |
| Intercepts mean | 0.426 | 0.426 | 0.547 | 0.358 | 0.503 | 1 | 22150 | 16917 |
| Intercept SD | 0.438 | 0.456 | 0.197 | 0.089 | 0.738 | 1 | 3688 | 6360 |
| **Predictors** | | | | | | | | |
| Slow wave | -0.388 | -0.386 | 0.237 | -0.765 | -0.008 | 1 | 34680 | 18981 |
| Fast wave | -0.781 | -0.777 | 0.259 | -1.197 | -0.370 | 1 | 41390 | 18324 |
| Large claw | 0.394 | 0.395 | 0.221 | 0.033 | 0.736 | 1 | 25805 | 18856 |
| Temperature | 0.447 | 0.447 | 0.527 | 0.405 | 0.490 | 1 | 25872 | 18631 |
| Carapace width | -0.150 | -0.148 | 0.160 | -0.394 | 0.114 | 1 | 21376 | 17198 |
| Claw:carapace ratio | -0.083 | -0.082 | 0.168 | -0.355 | 0.179 | 1 | 20397 | 16490 |
| Time relative to low tide | -0.001 | 0.000 | 0.118 | -0.179 | 0.198 | 1 | 25804 | 18517 |
| Treatment order | -0.360 | -0.359 | 0.221 | -0.709 | -0.004 | 1 | 24736 | 19644 |
| Phi (Beta dist. shape) | 6.927 | 6.712 | 1.670 | 4.333 | 9.349 | 1 | 5345 | 12260 |

Table 5: Summary statistics for the linear Hurdle model results looking at how the robotic rival affects the likelihood of entering the burrow and the proportion of time spent in-burrow. Shows summary statistics of the posterior distributions, $\hat{R}$, and effective sample sizes for the mean of intercepts, standard deviation of intercepts and predictor effect sizes. This is shown for both the probabilities of entering the burrow and the proportion of time spent in-burrow.
